# Adolescents’ dietary habits and meal patterns influence school performance in the Northern Finland Birth Cohort 1986: mendelian randomisation study

**DOI:** 10.1101/2021.04.30.442179

**Authors:** Loukas Zagkos, Fotios Drenos, Pauline Emmett, Alexandra I. Blakemore, Tanja Nordstrom, Tuula Hurtig, Marjo-Riitta Jarvelin, Terence M. Dovey

## Abstract

Several observational studies indicate that dietary habits in children and adolescents are associated with school performance. These associations are heavily confounded by socio-economic characteristics, such as household income and parents’ educational attainment, amongst other factors. In this study, we report observational and causal effects of habitual diet on school performance, using individual level data for 9,220 adolescents in the Northern Finland Birth Cohort 1986. For this purpose, we derived principal components for the dietary variables, meal patterns and school performance variables. The observational study showed a significant association of consumption of foods that are high in fat, salt and sugar (HFSS) with poor performance in all school subjects, and an association of consumption of healthy foods and traditional foods with good school performance in general subjects, science and physical education (PE). Moreover, a positive association was observed between not skipping breakfast and good performance in all school subjects. Mendelian randomisation analysis confirmed a negative effect of HFSS on school performance in general/science subjects (−0.080, −0.128 to −0.033) and a positive effect of healthy food on school performance in general/science subjects (0.071, 0.024 to 0.119) and PE (0.065, 0.021 to 0.110). To conclude, we identified compelling evidence that HFSS foods and healthy foods were causally affecting school performance.

## INTRODUCTION

Diets that contain a high proportion of fruit, vegetables, cereals and olive oil, moderate amounts of fish, dairy and low amounts of saturated fat and meat have been associated with a wide variety of positive health benefits, including improved cognitive function in older adults (Loughrey et al., 2017) and adolescents (Tapia-Serrano et al., 2021). The most recent theories suggest that these well-balanced diets reduce inflammation and/or oxidative stress and protect levels of brain-derived neurotrophic factor (Frisardi et al., 2010). In contrast, diets defined by high intake of fat, salt and sugar increase the likelihood of insulin resistance and have been identified as a leading cause of cognitive dysfunction (Allen et al., 2004; Fu et al., 2017; Pilato et al., 2020). These studies raise the possibility that particular habitual diets can protect against cognitive decline and may also be important for optimal cognitive development. Such advantageous diets may lead to improved scholastic attainment in early life, particularly reading comprehension (Haapala et al., 2016).

Individual dietary markers that enhance optimal cognitive performance or protect against cognitive decline remain elusive, and it may be more useful to consider more general dietary patterns. The most promising individual nutritional markers related to improving cognitive function/efficiency and mental health have been supplementation with either vitamin D (Azzam et al., 2015; Focker et al., 2017; Przybelski & Binkley, 2007; Hajiluian et al., 2017) or omega-3 polyunsaturated fatty acids (PUFA) (Patrick & Ames, 2015; Richardson et al., 2012; Sinn et al., 2010). Recent large scale replication studies have indicated that nutritional supplementation with omega-3 (PUFA) does not alter executive functions (Montgomery et al., 2018), although the correlation between eating fish (a primary source of omega 3 PUFA) and educational attainment remains (Pilato et al., 2020; Teisen et al., 2020). Taken together, whole diet or food quality assessments are likely to reveal larger impacts on intellectual development, mental health, behavioural problems and educational performance in children than single markers. A recent comprehensive review of the literature suggests that modulation of single nutrients within a whole diet is unlikely to result in improvements in educational attainment or performance (Kadosh et al., 2021).

Identifying a nutrient or food that will improve educational attainment and cognitive performance is laudable, but data indicate that a poor diet, defined by high intakes of fat, salt and sugar made up from a combination of foods, is associated with poor cognitive performance (Naveed et al., 2020) and neurobiological deficits (Stadterman et al., 2020). Observational studies have concluded that diets defined by processed foods (Wiles et al., 2009) or skipping meals (Lien, 2007) were associated with behavioural problems (Øverby & Høigaard, 2012) and poor cognitive performance (Pilato et al., 2020). However, “correlation is not causation” and it is important to determine whether these poor dietary patterns directly cause the associations observed, or are simply acting as markers of other factors (such as SES, for example). Indeed, determining whether a well-balanced diet improves/maintains (or a poor diet decreases) educational performance is the frontier of this field of research and one of the principal aims of the current study.

Previous attempts to assess diet have investigated specific nutrients rather than employing whole diet and have often relied on associations between food frequency questionnaires and single time-point psychometric assessments. This approach restricts conclusions and causes difficulty in assessing the impact of confounding variables (Kadosh et al., 2021). Food preferences are, at least partially, genetically determined, with heritability believed to be ranging between 36%-58% (Pallister et al., 2015) and specific polymorphisms have been associated with self-reported diet patterns (Guénard et al., 2017). This suggests that the application of Mendelian randomisation (MR) to disentangle the complex relationships between habitual diet and educational performance is possible. For this approach, genetic markers are used as proxies for characteristics that are difficult to reliably control in population studies (Evans & Davey-Smith, 2015). Since genetic variation in offspring is allocated randomly at conception, based on the genetic variation of parents, and is independent of other characteristics of the family, as postulated in the 2nd law of Mendel, genetic variants can be used as though they were interventions in a randomised control trial (Burgess et al., 2017). This allows us to use MR approaches to examine causative relationships between associated variables.

The aim of the current study was to explore the causal impact of habitual diet on cognitive function using population data from the Northern Finland Birth Cohort. Previous research has suggested that individuals with a balanced diet would achieve better outcomes than individuals who habitually consume comparatively higher quantities of fat, salt and sugar food items. It will also assess whether diet affects all scholastic subjects equally, supporting a general improvement in educational attainment, rather than specific effects. Here, we present data that clarify whether or not dietary patterns are indeed causative factors in school performance.

## METHODS

### Study sample

Northern Finland Birth Cohort 1986 (NFBC 86’) is a prospective cohort, which included mothers in the two northernmost provinces on Finland (Oulu and Lapland) with an expected date of birth between 1.7.1985 and 30.6.1986 (Järvelin M-R et al., 1993). Altogether 9,479 children were born into the cohort, 9,432 of them live-born. In addition to the data on delivery, follow-ups have been conducted at four different time points i.e. at 1, 7-8, 15-16 and 33-35 years of age. Collected data include prenatal and early life measurements, information on motor, social, psychological and mental development in childhood. In adulthood data has been collected on social background, lifestyle, medication, diagnosed diseases, organ-specific and psychiatric symptoms, workload and occupational health, economy, personal traits, functioning, quality of life, and family history of diseases. More detailed description can be found at https://www.oulu.fi/nfbc/.

### Dietary data

The adolescents’ 16-year follow-up questionnaire contained a set of questions on the frequency of consumption of a wide variety of foods. For example, they were asked how many slices of bread they had in a day, and how often they had eaten uncooked vegetables the previous week. Moreover, they were asked about the frequency of having consumed specific foods during the previous six months. All dietary data were numerically transformed into ‘times consumed per week’, in order to apply quantitative meaning to the diet variables. In the same questionnaire, the adolescents were asked whether they usually had breakfast/ lunch/ dinner on weekdays and on weekends, in a Yes/No question. To assess their meal patterns, variables were transformed into ‘times had breakfast/ lunch/ dinner per week’. Collecting data in this manner and from this age group has been found to be valid and reliable with a long history of effective assessment of diet (Rockett et al., 1997; Vilela et al., 2018; Willet et al., 1987).

### School performance

In the 16-year follow-up study, adolescents were asked to report how well they were doing in various school subjects: Finnish language, general subjects, such as history and religious studies, mathematics, biology, physics, chemistry, art and physical education, compared to other pupils of their age. The question had four levels of possible answers: Better than average, average, worse than average and really badly.

### Genetic data

Genetic information was available for 3,834 individuals. Quality control was performed as suggested by the Northern Finland Birth Cohort documentation. We excluded 26 samples from the analysis, determined to be outliers for ‘missingness’ and ‘heterozygosity’ and 4 samples due to sex mismatch. In addition, another 24 samples were removed due to relatedness and 37 samples as duplicates (checked by the ID numbers). After filtering, a total of 3,743 individuals remained for analysis. In total, 928,272 autosomal single nucleotide polymorphisms (SNPs) were available prior to imputation. SNPs with a Hardy-Weinberg equilibrium p-value threshold smaller than 10^−4^ were removed from the analysis. The total number of genotyped SNPs that passed the quality control was 889,119. Imputation was performed using the 1000 Genomes phase 3 reference panel. From 81,571,831 imputed and directly genotyped SNPs, the variants with an imputation info score lower than 0.9 were excluded. Only variants with minor allele frequency greater than 0.01 were included. After filtering, 11,009,294 variants remained.

### Imputation of missing values

Individuals with up to 6 missing dietary values (out of 38) and 1 missing school performance value (out of 6) were excluded from the analysis. To address missing data and gain statistical power on the observational calculations a multivariate imputation by chained equations (MICE) was created. The MICE algorithm creates multiple imputations – replacement values – for multivariate missing data. Creating multiple imputations, as opposed to single imputations, accounts for the statistical uncertainty in the imputations and maintains an unbiased sample variance. For both datasets, the predictive mean matching imputation method with 5 multiple imputations and 50 iterations on each step for high accuracy was performed.

### Statistical analysis

#### Statistical software

Analysis was conducted in R version 4.0.2 (R Core Team, 2020) and figures were produced using the R package “forestplot” (Gordon & Lumley, 2020).

#### Principal component analysis

Principal component analysis (PCA) with varimax rotation was performed separately for dietary variables (33), meal patterns (6) and school performance variables (6). The purpose was to reduce the dimensions of the dietary dataset but also to increase the interpretability of all datasets. For this analysis we used the Bartlett factor score method, which is based on maximum likelihood estimates and provides unbiased estimates of the true factor scores (Hershberger, 2005). Factor loadings were calculated for each variable, denoting the amount of variable information described by each factor. The number of factors that best represents the data was determined by performing a scree test (Mardia et al., 1979). Foods and meal patterns with loadings above 0.30 on a factor were considered to have a strong association with that factor and were the most informative in describing the dietary factors (Northstone & Emmett, 2008). School performance variables with factor loadings greater than 0.4 were considered most representative of these factors.

#### Multivariable regression of observational data

In the observational analysis, multivariable linear regression to estimate the association of diet with school performance was undertaken. The NFBC 86’ data were used to adjust the analysis for potential confounders: sex, household income, family financial status, mother and father education. Regressions for each school performance factor against every dietary factor and meal pattern factor were then completed.

#### Genome wide association analysis

Under the additive genetic model, we tested for associations between 11,009,294 genetic variants that passed quality control and 5 dietary and 3 school performance principal components for up to 3,743 individuals in the NFBC 86’. The association of each genetic variant was tested using a regression model adjusting for age and the first 4 genetic principal components to control for population structure. We used the FUMA web application (Watanabe et al., 2017) to generate Manhattan and Quintile-Quintile plots and to annotate genetic associations. Moreover, the Genotype-Tissue Expression (GTEx) project database was used to assess the expression of the identified genes in various human tissues. Gene expression analysis from GTEx is integrated in FUMA.

#### One-sample mendelian randomisation

Two-stage least squares (2SLS) regression analysis was performed in the NFBC 86’ to investigate the causal relationship between diet and school performance. For each dietary exposure, independent variants (r^2^<0.01) were used as genetic instruments for the analysis at a P-value threshold of 10^−4^. In the first step of the analysis, each dietary factor was regressed against the genetic instruments and the measured confounders: sex, household income, family financial status, mother’s, and father’s education level. In the second step of the analysis, each school performance factor was regressed against the fitted values from the first step of the analysis.

To assess the validity of the MR assumptions, one-sample mendelian randomisation sensitivity analysis was performed. To test whether the genetic variants used in the analysis were associated with the potential confounders, the confounders were regressed against the exposure of significant genetic variants. SNPs with the evidence of association with the confounders were then removed and the two-stage least squares regression analysis was repeated to investigate whether this significantly altered the regression results. The robustness of the results to violations was tested by applying two sample MR methods on the data, including MR–Egger (Bowden et al., 2015), MR weighted median (Bowden et al., 2016) and MR weighted mode (Hartwig et al., 2017) analyses.

## RESULTS

The demographic characteristics of the sample can be seen in **Table 1**.

**Table 1.** Characteristics of the sample used in the analysis.

|  |  |
| --- | --- |
| % Sex |  |
| Male | 50.6 |
| Female | 49.4 |
| % Mother education |  |
| Less than 9 years | 4.4 |
| Comprehensive school education | 60.3 |
| Matriculation examination | 35.3 |
| % Family financial status |  |
| Many problems | 2.2 |
| Some problems | 19.5 |
| Quite good status | 66.9 |
| Very good status | 11.4 |
| % Father education |  |
| Less than 9 years | 8.5 |
| Comprehensive school education | 71.6 |
| Matriculation examination | 19.9 |

### Principal component analysis

In order to identify food clusters that share common variation and can be explained by the same principal component, dietary data on 5,337 adolescents were included in the analysis.

For the diet analysis, the first three principal components were determined sufficient to describe the data, explaining 21% of total variance in the dietary dataset. Little variance was explained by the rest of the principal components. The first component was characterised by foods high in fat, salt and sugar and thus named “*HFSS*” food. In the second component, chicken, fish, rice and pasta loaded highly, so we refer to this as “*healthier*” food choices. The third component was characterised by foods that constitute traditional Finnish meals, such as different kind of meats, boiled potatoes, baked goods. Therefore, factor three was named “*traditional*” food. Details of the groups can be seen in **Table 2**.

**Table 2.** Principal factors of dietary variables. Foods with loadings above 0.3 are denoted with x.

| **Variables** | **Factor 1** | **Factor 2** | **Factor 3** |
| --- | --- | --- | --- |
| Glasses of milk | - | - | - |
| Glasses of sour milk | - | - | - |
| Glasses of other milk products | - | - | - |
| Slices of dark bread | - | - | - |
| Slices of full wheat or oatmeal bread | - | - | - |
| White bread | - | - | - |
| Uncooked veg | - | - | - |
| Uncooked fruit | - | - | - |
| Berries | - | - | - |
| Cakes and cookies | - | - | **x** |
| Porridge | - | - | - |
| Cold breakfast | - | - | - |
| Cheese | - | - | - |
| Ice-cream | **x** | - | - |
| Boiled potatoes | - | - | **x** |
| Fried potatoes or French fries | **x** | - | - |
| Rice | - | **x** | - |
| Pasta | **x** | **x** | - |
| Fish | - | **x** | - |
| Chicken | - | **x** | - |
| Sausages | **x** | - | **x** |
| Cold meats | - | - | **x** |
| Meat dishes | - | - | **x** |
| Ground beef | - | - | **x** |
| Eggs | - | - | - |
| Soft drinks with sugar | **x** | - | - |
| Soft drinks without sugar | - | - | - |
| Chocolate | **x** | - | - |
| Sweets | **x** | - | - |
| Ready-to-eat food | **x** | - | - |
| Salad dressing | - | - | - |
| Hamburgers, pizza | **x** | - | - |
| Potato crisps | **x** | - | - |

Similarly, principal component analysis on the six meal patterns revealed two principal factors accounting for 15% of total variance in meal pattern variables. The cumulative explained variance did not significantly change when considering a third component. Breakfast was the only variable that characterised the first principal component, whereas evening snacks and snacks between meals had large loadings in the second principal component (**Table 3**). As a result, principal component one is referred to as “*breakfast*” and principal component two as “*snacks*”.

**Table 3.** Principal factors of meal. Variables with loadings above 0.3 are denoted with x.

| **Variables** | **Factor 1** | **Factor 2** |
| --- | --- | --- |
| Breakfast | **x** | - |
| Lunch | - | - |
| Dinner | - | - |
| Evening snack | - | **x** |
| Snack between meals | - | **x** |
| Devour food | - | - |

Following the same analysis for school performance, the first three principal components explained 60% of total variance of the variables. The rest did not explain a significant proportion. The first factor, which explains 24% of the total variance, is described by general subjects (history and religious studies), mathematics and physical sciences - biology, physics and chemistry (**Table 4**). Finnish language had a 0.96 loading on the second factor, which accounts for 20% of the total variance. Physical education has a loading of 0.99 on the third factor, explaining 16% of total variance. Consequently, factors 1, 2 and 3 were named “*general/science*”, “*Finnish*” and “*Physical Education* (*PE*)”, respectively. **Supplementary tables 1**, **2** and **3** show all factor loadings for diet, meal patterns and school performance variables.

**Table 4.** Principal factors of school performance variables. Variables with loadings above 0.4 are denoted with x.

| **Variables** | **Factor 1** | **Factor 2** | **Factor 3** |
| --- | --- | --- | --- |
| Finnish language | - | **x** | - |
| General subjects | **x** | - | - |
| Mathematics | **x** | - | - |
| Biology, Physics, Chemistry | **x** | - | - |
| Art | - | - | - |
| Physical Education | - | - | **x** |

### Observational association of diet on school performance

Each school performance factor was regressed against every dietary factor and meal pattern factor to identify associations between diet and school performance, adjusting for measured confounders: sex, household income, family financial status, mother’s and father’s education. **Figure 1** shows that *HFSS* foods were negatively associated with school performance in *general/science* and *Finnish* with an effect size of −0.159 (95% confidence interval −0.190 to −0.127) and −0.069 (−0.099 to −0.040), respectively. There was no evidence of association between *HFSS* food and *PE*.

**Figure 1.**
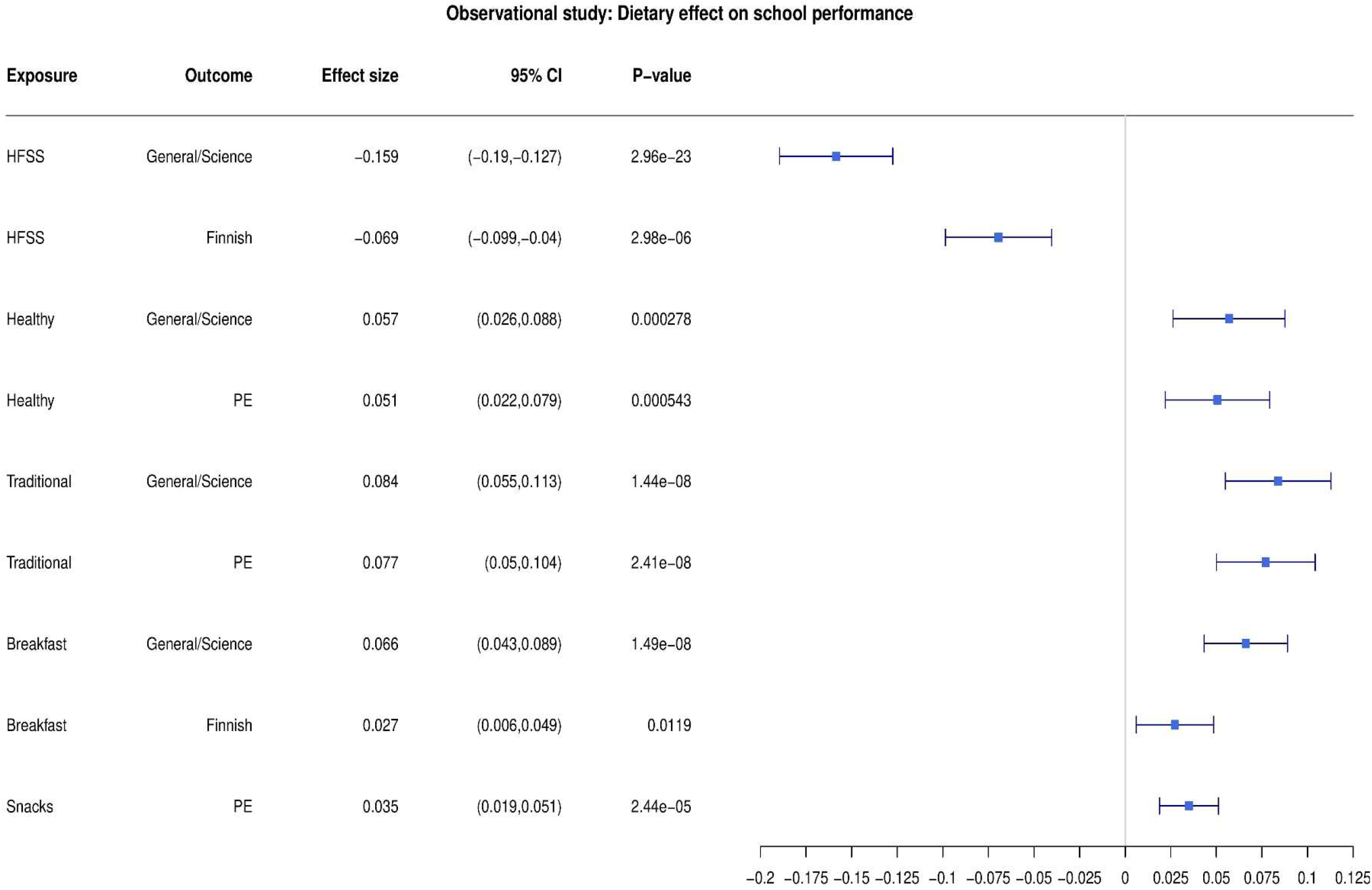
Results of the observational study: the effect of an additional weekly meal in dietary exposures on school performance outcomes

Healthier eating was positively associated with better school performance in *general/science* (0.057, 0.026 to 0.088) and *PE* (0.051, 0.022 to 0.079), as was *traditional* meal eating with school performance in *general/science* (0.084, 0.055 to 0.113) and *PE* (0.077, 0.050 to 0.104). No significant association was found between *healthier* eating or *traditional* eating with school performance in *Finnish*.

Eating *breakfast* was positively associated with school performance in *general/science* (0.066, 0.034 to 0.089) and *Finnish* (0.027, 0.006 to 0.049). In addition, there was evidence that eating *snacks* was positively associated with good performance in *PE* (0.035, 0.019 to 0.051). No evidence of association was found between *snack* consumption and school performance neither in *general/science* and *Finnish* language, nor between eating *breakfast* and school performance in *Finnish* language and *PE*.

### Genome wide association results

Genome wide association with the diet patterns identified 403 SNPs associated with *HFSS*, 282 with *healthier* eating, 207 with *traditional* meal eating, 154 with eating *breakfast*, and 191 with eating *snacks* (**Supplementary table 4** and **Supplementary figures 1**-**5**). The p-value selected as a cut-off in this work was 10^−4^, as it was the threshold that provided robust signals for all dietary GWAS and moreover, took into account the relatively small sample size of NFBC 86’ compared to existing GWAS of large cohort studies. Integration of association results with GTEx gene expression across 54 tissue types through FUMA (Watanabe et al., 2017), showed that all five dietary patterns were associated with genes up-regulated in the brain with borderline significance. (**Supplementary figures 6**-**10**). SNPs associated with *HFSS* had previously been reported as associated with heel bone density (Kim, 2018), blood cell measures (Astle et al., 2016), autoinflammatory problems (Zhu et al., 2018; Paternoster et al., 2015) and cardiometabolic relevant traits such as pulse pressure and high-density lipoprotein (HDLc) cholesterol levels (Kanai et al., 2018; Evangelou et al., 2018). SNPs for *healthier* eating had previously been reported as associated with physical activity (Doherty et al., 2018), being a morning person (Jones et al., 2019), and loneliness (Day et al., 2018). SNPs associated with a *traditional* diet, eating *breakfast* and eating *snacks* have not been previously reported as associated with traits relevant to diet.

### MR effects of diet on school performance

A one-sample MR was performed on the significant associations between diet and school performance. Independent genetic variants identified in the GWAS as associated with the dietary exposures were used as instruments. **Figure 2** summarises the results of two-stage least squares regression. *HFSS* had a negative effect on school performance in *general/science*, with an effect size of −0.08 (−0.128, −0.033). There was no evidence of a causal MR relationship between *HFSS* and school performance in *Finnish*. The MR results were in agreement with the observational study, showing a positive effect of *healthier* food consumption (primarily chicken, fish, rice and pasta) on school performance in *general/science* (0.071, 0.024 to 0.119) and *PE* (0.065, 0.021 to 0.110). The analysis also confirmed that *traditional* food consumption was associated through MR with better educational outcomes in *general/science* (0.066, 0.017 to 0.116) and *PE* (0.116, 0.07 to 0.162). Eating *breakfast* had a positive effect on school performance in *general/science* with an effect size of 0.126 (0.081,0.171) and a very strong p-value of 3.4×10^−8^. There was not enough evidence to support the observed positive association of *eating breakfast* with school performance in *Finnish* language. Lastly, eating *snacks* was associated with a positive MR effect in *PE* performance (0.046, 0.016 to 0.075).

**Figure 2.**
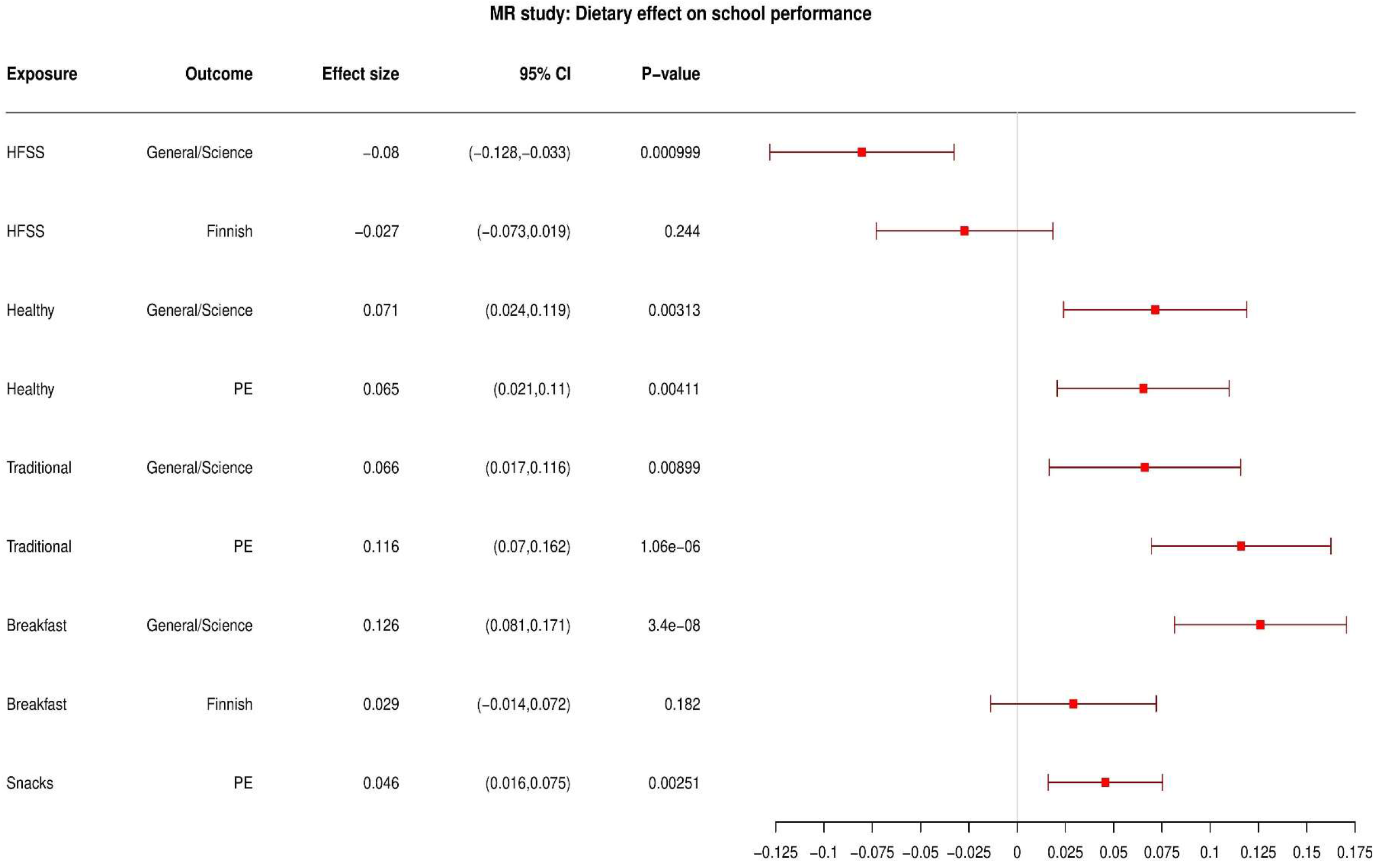
Results of one-sample mendelian randomisation: the effect of an additional weekly meal in dietary exposures on school performance outcomes.

### Sensitivity analyses

**Supplementary table 5** shows the genetic variants associated with at least one measured confounder below the threshold of p=10^−4^. The forest plot in **Supplementary figure 11** shows that the excluded variants, which were associated with the measured potential confounders, do not influence the significance of the causal associations between diet and school performance. The robustness of our results was tested through the use of other MR methods, including MR-Egger, weighted median and weighted mode MR, that are less sensitive to small violations of our assumptions, such as pleiotropy. The results in **Supplementary tables 6**, **7**, **8** and **9** show no evidence of pleiotropic effects in the variants involved in the analysis or a substantial change of our results.

## DISCUSSION

The purpose of this work was to explore the causal effect of habitual diet on school performance in adolescents in the NFBC 86’. Our observational and genetic findings support the concept that dietary patterns causally influence performance in various school subjects. More specifically, in the observational part of this study, significant negative associations between HFSS food and school performance, and positive associations between healthy eating, traditional meal eating, eating breakfast, and having snacks was found with school performance in various subjects. These outcomes were confirmed by the MR analysis findings. Strong causal effects of the dietary factors on school performance in various subjects, except for performance in Finnish language, were identified.

The traditional Finnish diet, up until the 1960’s, was typified by a diet high in fat and saturated fat (Prättälä, 2003). With increased globalisation and socioeconomic changes, the Finnish diet has progressively altered in line with European health-related drives and lifestyles. Increases in vegetables and low-fat alternative products have been observed (Pietinen et al., 2001). In Finland, as in all developed countries, there has been an increased demand for plant-based products and a move away from traditional meat and dairy food choices (Huan-Niemi et al., 2020). Primary motivators for this change in behaviour relate to climate change (Röös et al. 2018) and perceived health benefits (Willet et al., 2019).

The current study supports and extends previous findings. Habitual diets rich in fish and low saturated fat meats enhance educational attainment (Loughrey et al., 2017; Tapia-Serrano et al., 2021), while consuming HFSS foods, such as fried potatoes, sausages, soft drinks with sugar, chocolate, sweets, ready-to-eat meals, hamburgers, pizza, potato crisps and ice-cream, appears to be detrimental to educational attainment (Naveed et al., 2020). Timing of eating and specifically skipping breakfast (Lien, 2007) were also confirmed as detrimental within this population. Among these dietary patterns, it should be noted that fruit and vegetable intakes did not represent a significant component of any of the main patterns found. This outcome is aligned with a novel hypothesis that the impact of diet on educational attainment is associated with protein and carbohydrate macronutrient intakes rather than with micronutrient intakes (Jansen et al., 2020; Teisen et al., 2020). This suggestion needs to be investigated further with analysis of blood metabolites against markers of diet and their impact on cognitive performance.

MR analysis showed that an additional weekly HFSS meal consumed causes an 8% decrease in school performance in general/science subjects. Moreover, an additional healthy meal per week was identified to cause a 7.1% increase in school performance in general/science subjects and a 6.5% increase in PE. Interestingly, a 6.6% raise in school performance in general/science subjects and a 11.6% increase in PE was found to be caused by an additional traditional Finnish meal consumed per week. Eating breakfast daily was identified to have the strongest causal effect on school performance in general/science subjects in our analysis. A one unit increase in having breakfast can cause a 12.6% increase in school performance in general/science subjects. Last, a unit increase in having snacks between meals and evening snacks was identified to cause a 4.6% increase in school performance in PE.

Information exploring the quality of the food eaten is still in its infancy, but the interpretation of our data supports the premise that dietary quality may play an influential role (Kadosh et al., 2021). Focus on assessing the dose dependent effects of healthier food intake in children and its impact on education poses few ethical concerns than exploring the detriment incurred by consuming HFSS foods. However, these data assert that it is within the adolescents with HFSS diets and skipping breakfast that the most educational attainment gains could be achieved. Researchers should also be aware that the relative impact of diet on educational attainment explains, at best, 13% of the overall variation in educational attainment. Most variables within this study explained 5% of the variation. Therefore, focus on diet should be tempered by realistic expectations of the potential impact their modulation could have.

Some limitations are evident in this study. Diet is a cultural characteristic and the associations observed here may not necessarily apply to other cultural backgrounds; although with increased globalisation and equity in development, diet does appear to be homogenising. Furthermore, our approach of linking diet to genetics and, by extension, to biological changes, makes it more likely that these findings will be more widely valid. The genetics of diet, especially at the adolescent stage of life, are still underexplored, which has prevented replication of the identified MR associations. Related to this point, the sample size used was large compared to other published studies but still modest in terms of power. Some of the instances where an MR association was not observed could be attributed to lack of statistical power.

To summarise, these analyses provide evidence, through Mendelian randomisation that diet may have a causal effect on school performance. Thus, this work suggests that dietary interventions might be able to increase pupils’ educational attainment. However, this should be considered as only one part of the environment relevant to education. The identified causal effects are part of a wider and more complex causal network where additional interventions could reduce, or amplify, what was observed in this population. Considering the same questions in different educational and cultural settings would aid in understanding these relationships. Mapping the biological processes underpinning these findings will be integral to uncovering unbiased outcomes. Finally, providing, healthy and nutritionally balanced school meals to all children may improve not only their physical health, but also extend the benefits to education outcomes; however, eliminating HFSS and not skipping breakfast will have a larger effect on educational outcomes than providing healthier alternatives.

## ACKNOWLEDGEMENTS

We thank all cohort members and researchers who have participated in the NFBC study. We also wish to acknowledge the work of the NFBC project centre.

## SUPPLEMENTARY MATERIAL

**Supplementary table 1.** Factor loadings for all dietary variables in principal component analysis.

| Variables | Factor 1 | Factor 2 | Factor 3 |
| --- | --- | --- | --- |
| Glasses of milk | 0.02 | -0.02 | 0.08 |
| Glasses of sour milk | 0.01 | 0.11 | 0.06 |
| Glasses of other milk products | 0.04 | 0.09 | 0.13 |
| Slices of dark bread | -0.05 | 0.07 | 0.12 |
| Slices of full wheat or oatmeal bread | -0.01 | 0.08 | 0.12 |
| White bread | 0.22 | 0.05 | 0.01 |
| Uncooked veg | -0.15 | 0.27 | 0.10 |
| Uncooked fruit | -0.08 | 0.28 | 0.08 |
| Berries | -0.05 | 0.22 | 0.21 |
| Cakes and cookies | 0.06 | -0.07 | 0.30 |
| Porridge | -0.10 | 0.07 | 0.24 |
| Cold breakfast | 0.06 | 0.10 | 0.11 |
| Cheese | -0.01 | 0.08 | 0.25 |
| Ice-cream | 0.36 | 0.05 | 0.17 |
| Boiled potatoes | -0.01 | 0.09 | 0.50 |
| Fried potatoes or French fries | 0.50 | 0.26 | 0.05 |
| Rice | 0.22 | 0.68 | -0.05 |
| Pasta | 0.30 | 0.55 | 0.05 |
| Fish | 0.17 | 0.60 | 0.15 |
| Chicken | 0.23 | 0.69 | 0.07 |
| Sausages | 0.33 | 0.20 | 0.36 |
| Cold meats | 0.17 | 0.03 | 0.42 |
| Meat dishes | 0.21 | 0.21 | 0.59 |
| Ground beef | 0.25 | 0.20 | 0.54 |
| Eggs | 0.25 | 0.26 | 0.28 |
| Soft drinks with sugar | 0.51 | -0.04 | 0.10 |
| Soft drinks without sugar | 0.28 | 0.11 | 0.03 |
| Chocolate | 0.56 | 0.00 | 0.05 |
| Sweets | 0.55 | -0.07 | 0.05 |
| Ready-to-eat food | 0.45 | 0.09 | -0.01 |
| Salad dressing | 0.23 | 0.21 | 0.11 |
| Hamburgers, pizza | 0.65 | -0.01 | -0.02 |
| Potato crisps | 0.69 | 0.01 | 0.00 |

**Supplementary table 2.** Factor loadings for meal patterns.

| Variables | Factor 1 | Factor 2 |
| --- | --- | --- |
| Breakfast | 0.69 | 0.27 |
| Lunch | -0.04 | 0.28 |
| Dinner | 0.14 | -0.02 |
| Evening snack | 0.13 | 0.31 |
| Snack between meals | -0.01 | 0.32 |
| Devour food | -0.11 | 0.11 |

**Supplementary table 3.** Factor loadings for school performance variables.

| Variables | Factor 1 | Factor 2 | Factor 3 |
| --- | --- | --- | --- |
| Finnish language | 0.26 | 0.96 | -0.03 |
| General subjects | 0.47 | 0.37 | 0.08 |
| Mathematics | 0.57 | 0.23 | 0.06 |
| Biology, Physics, Chemistry | 0.90 | 0.16 | 0.05 |
| Art | 0.07 | 0.17 | 0.05 |
| Physical Education | 0.08 | 0.10 | 0.99 |

**Supplementary table 4.** Number of significant variants used as instruments for each variable in the one-sample mendelian randomisation setting.

| Trait | Number of variants |
| --- | --- |
| HFSS | 403 |
| healthy | 282 |
| traditional | 207 |
| breakfast | 154 |
| snacks | 191 |

**Supplementary figure 1.**
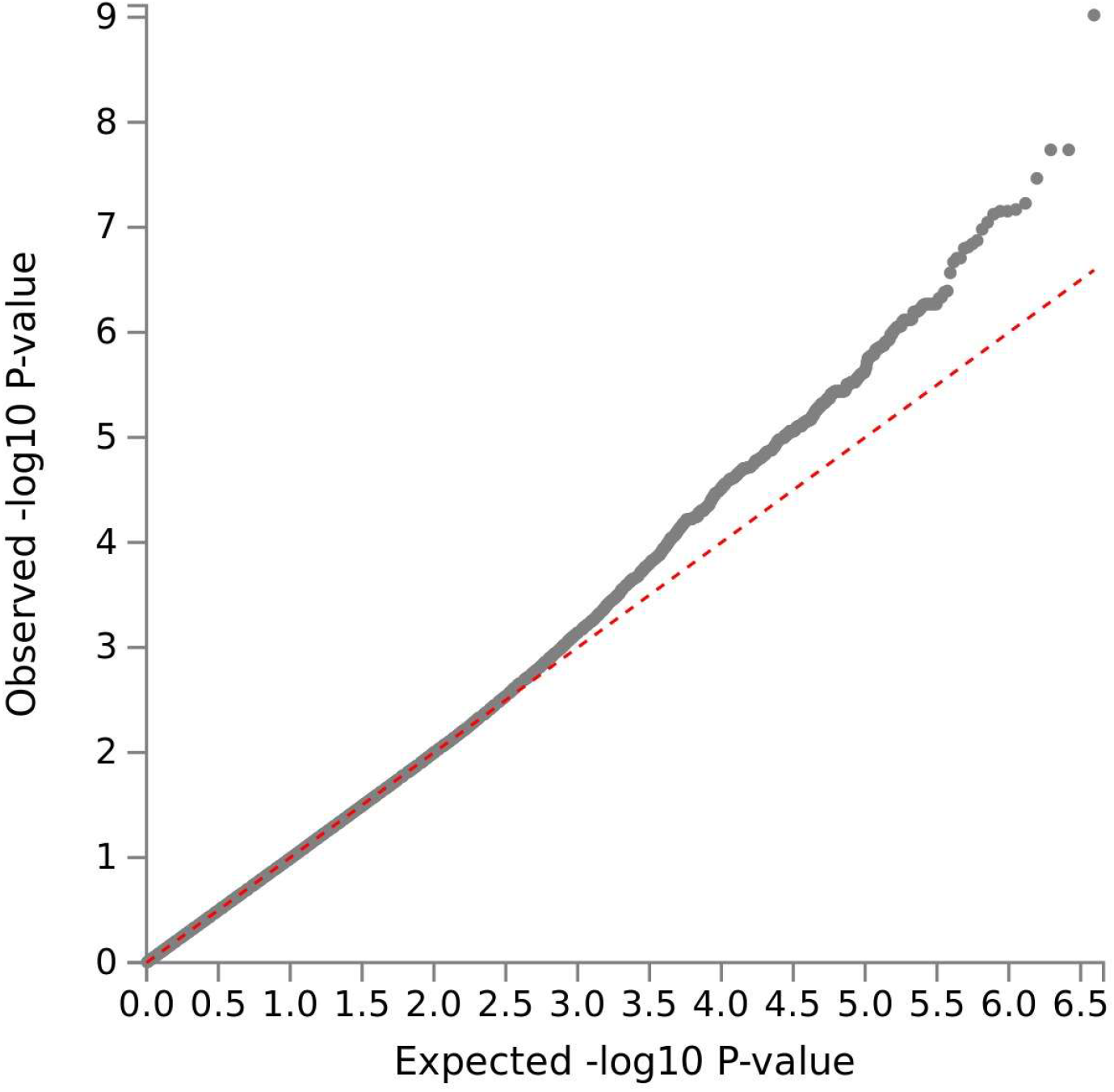
Q-Q plot of HFSS food consumption summary statistics, generated by FUMA.

**Supplementary figure 2.**
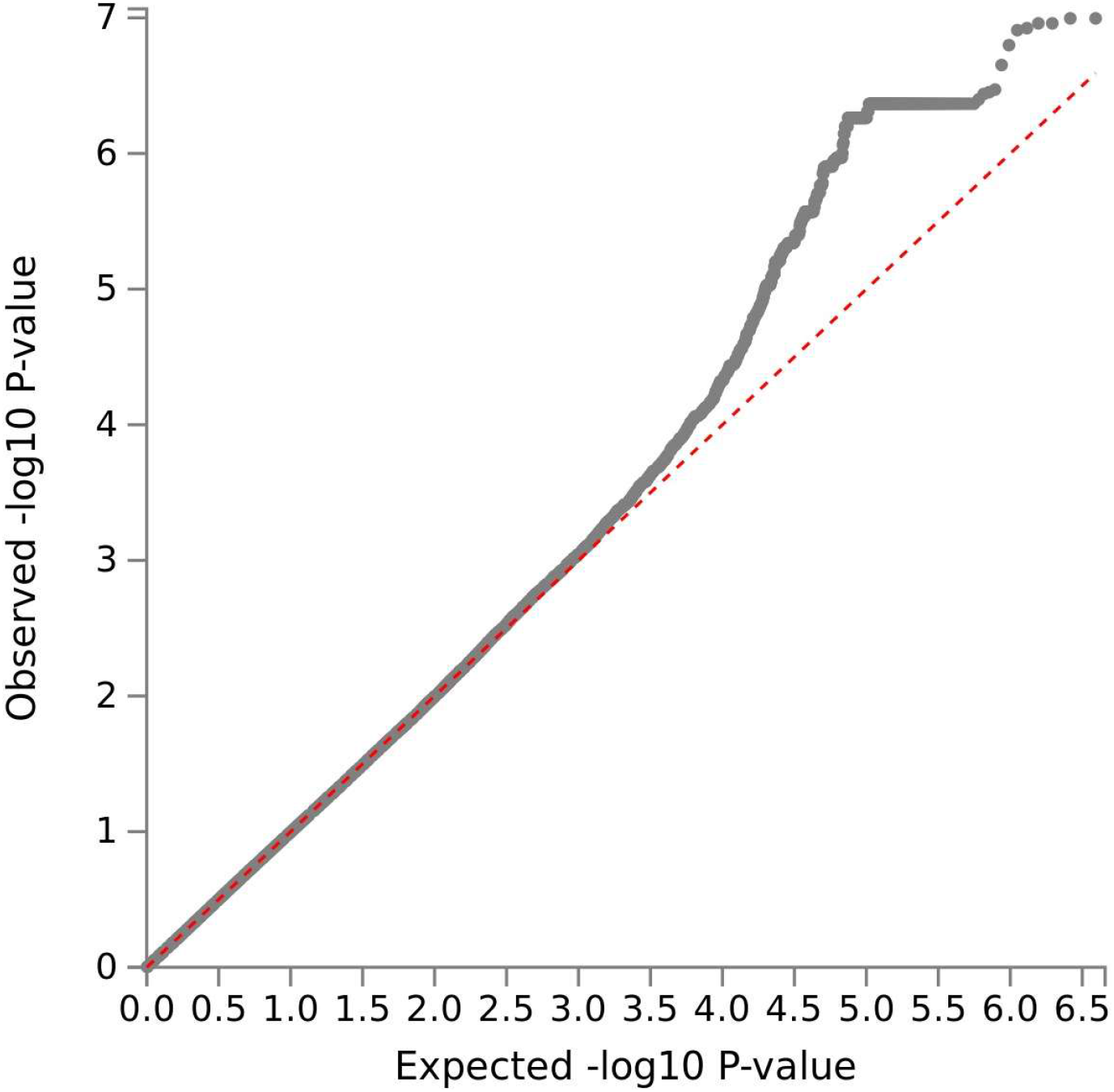
Q-Q plot of healthy eating summary statistics, generated by FUMA.

**Supplementary figure 3.**
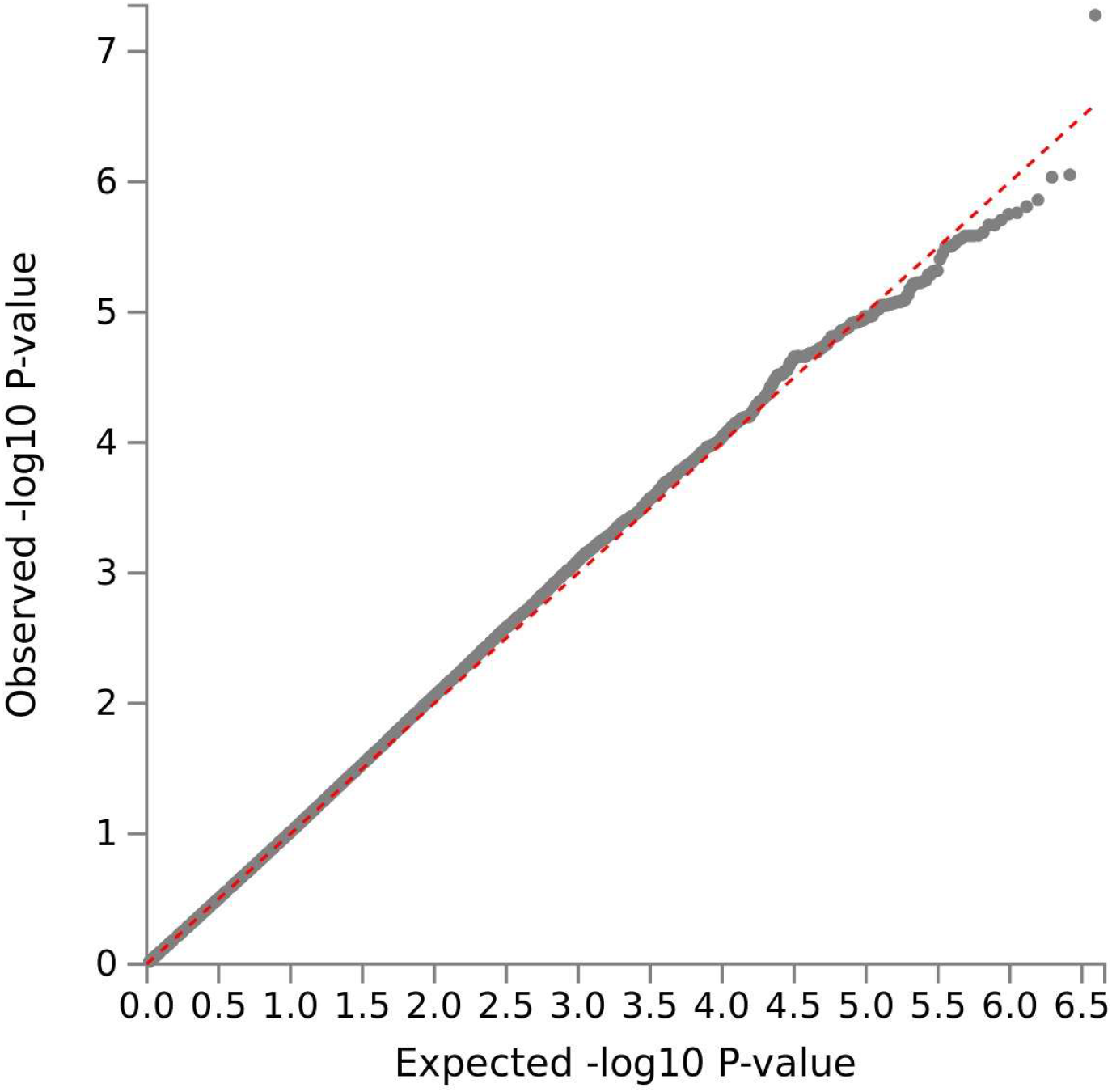
Q-Q plot of traditional meal consumption summary statistics, generated by FUMA.

**Supplementary figure 4.**
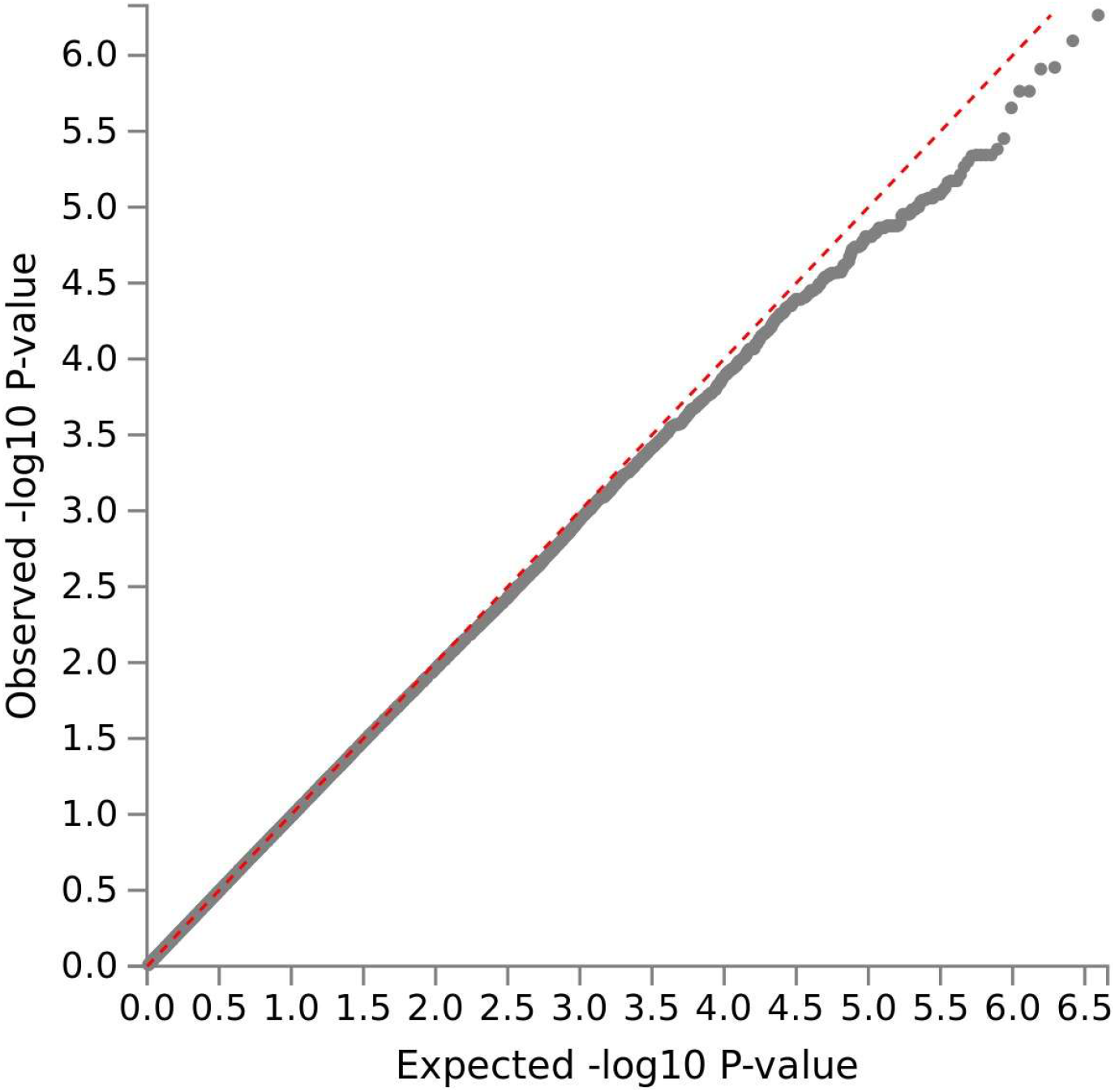
Q-Q plot of breakfast eating summary statistics, generated by FUMA.

**Supplementary figure 5.**
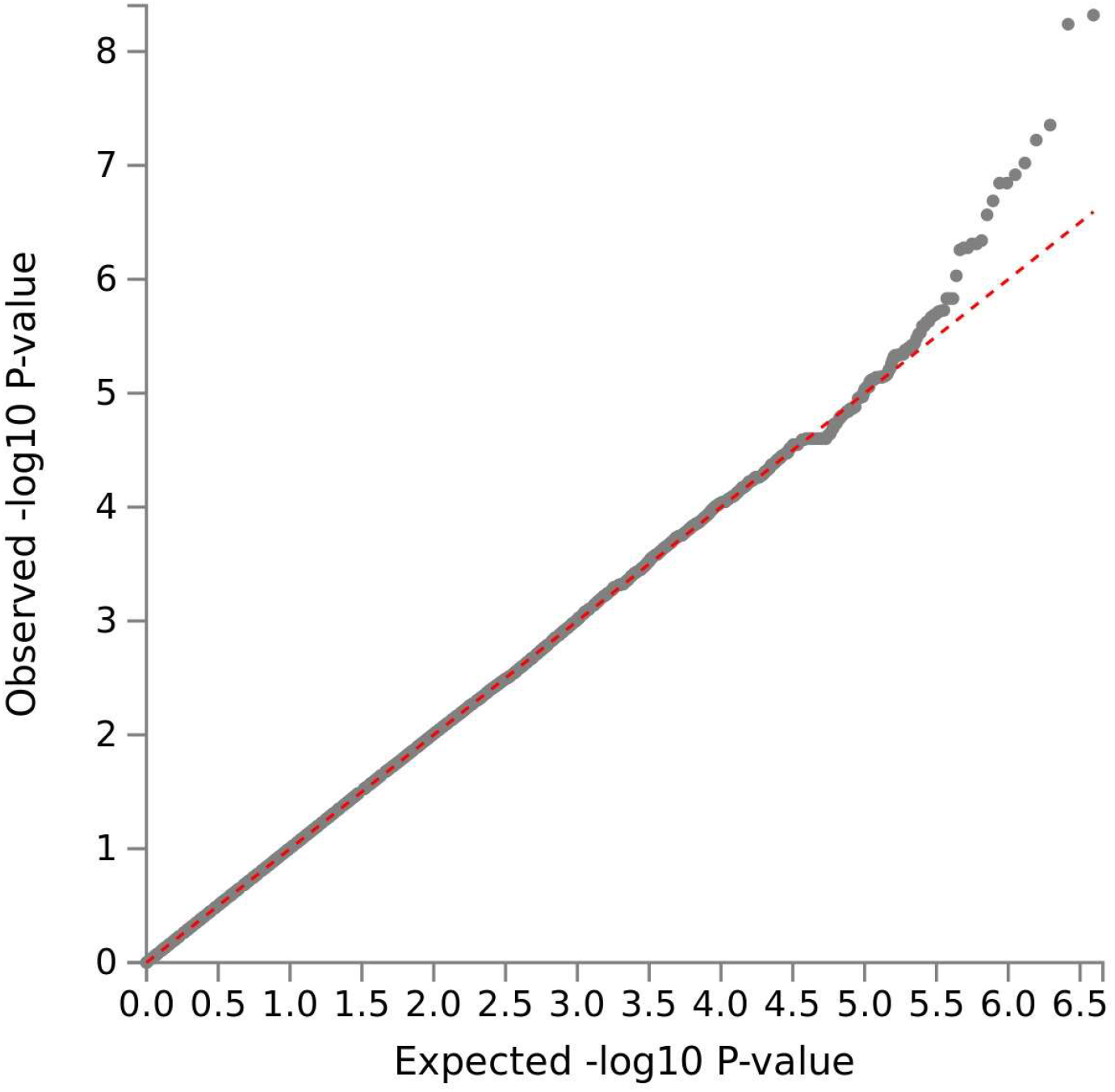
Q-Q plot of snack eating summary statistics, generated by FUMA.

**Supplementary figure 6.**
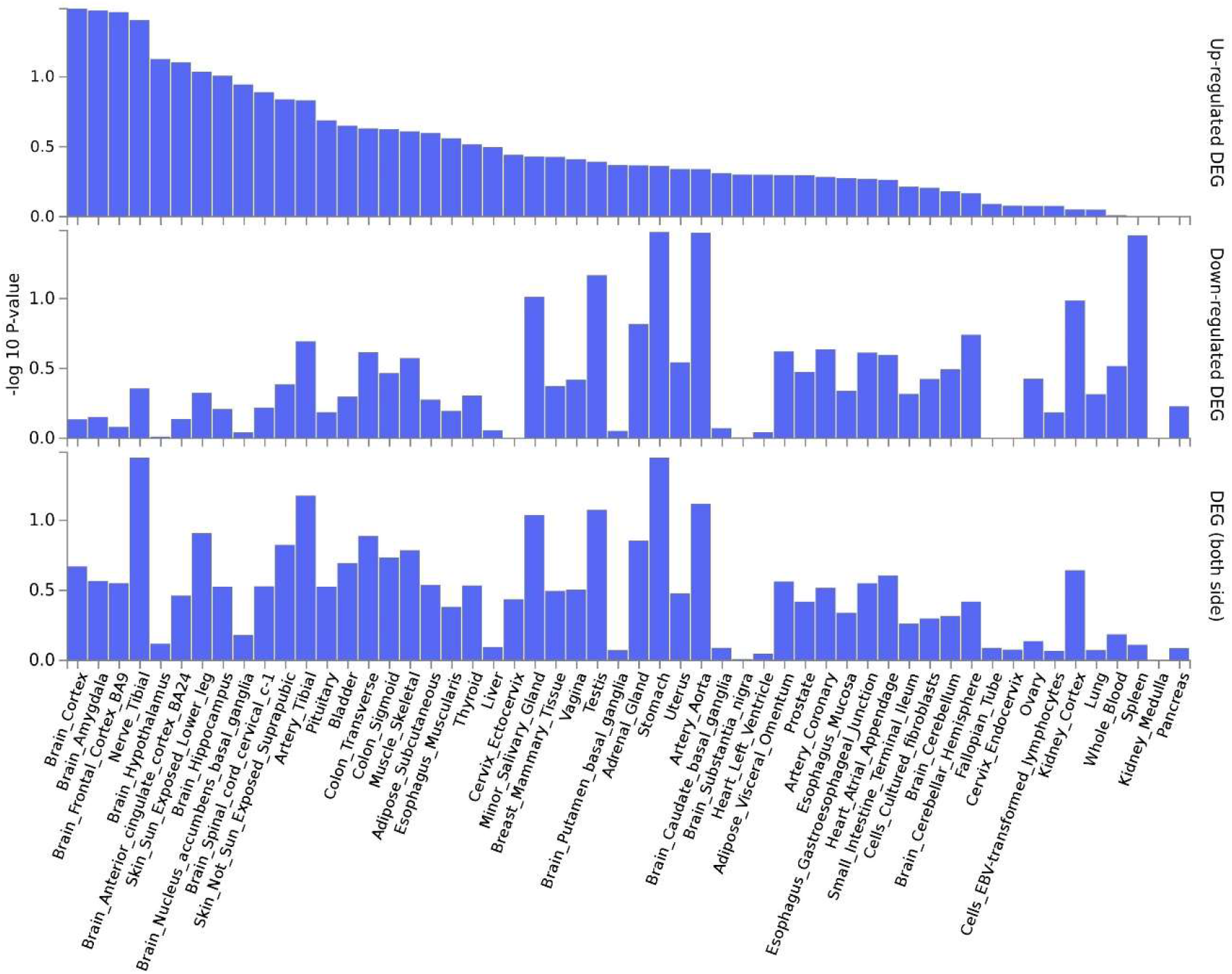
Up-regulated and down-regulated differentially expressed genes associated with HFSS food, generated by FUMA.

**Supplementary figure 7.**
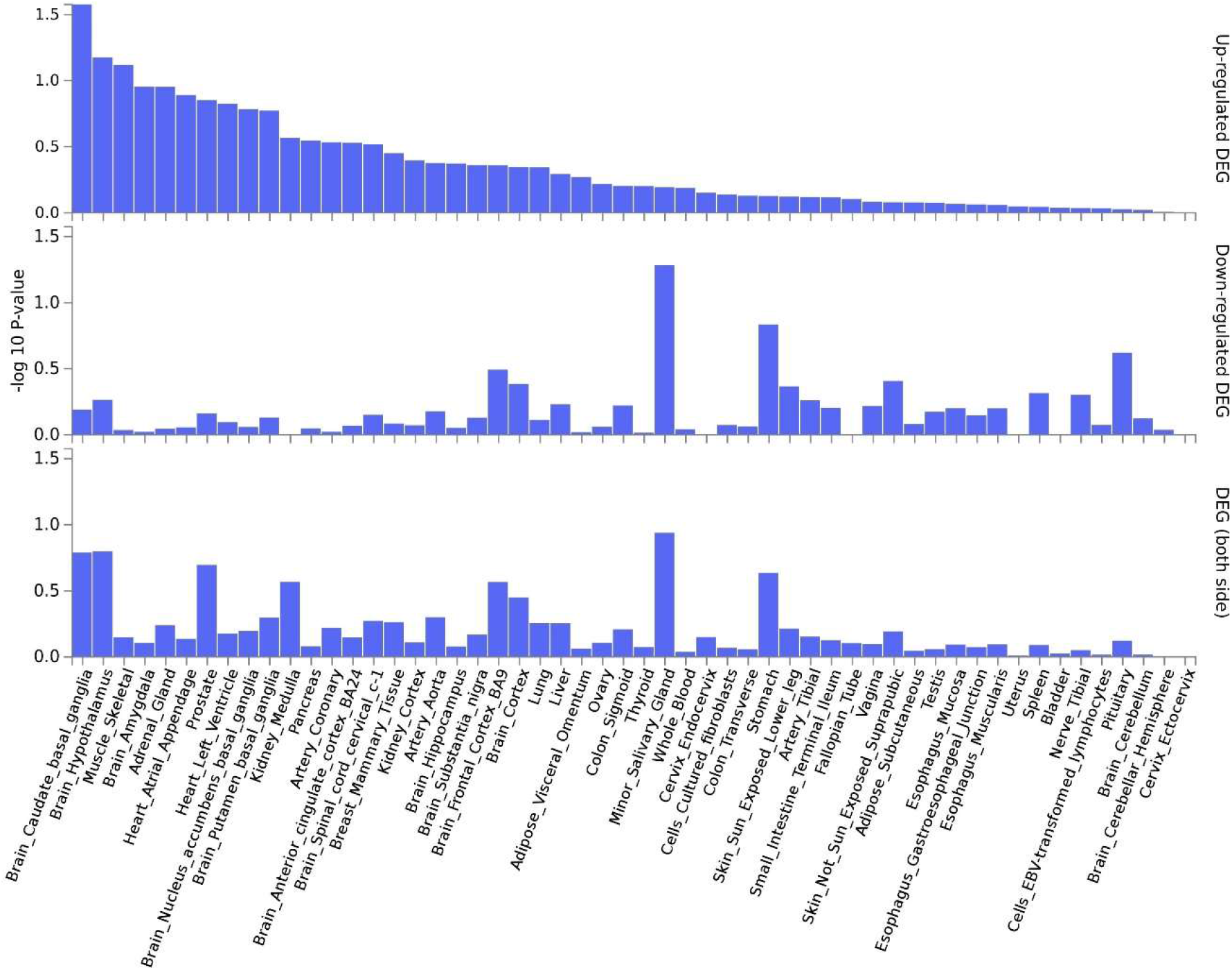
Up-regulated and down-regulated differentially expressed genes associated with healthy eating, generated by FUMA.

**Supplementary figure 8.**
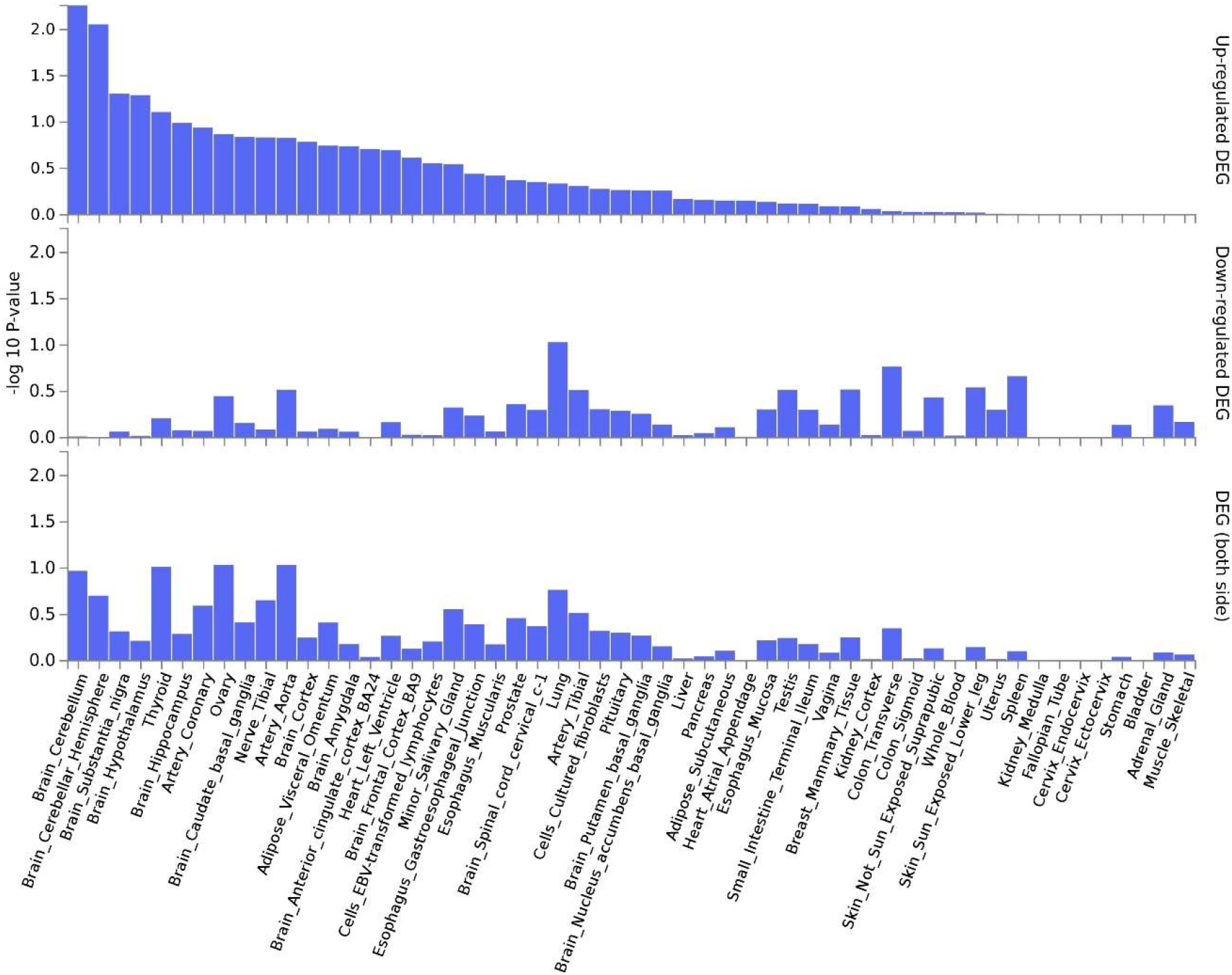
Up-regulated and down-regulated differentially expressed genes associated with traditional Finnish meals, generated by FUMA.

**Supplementary figure 9.**
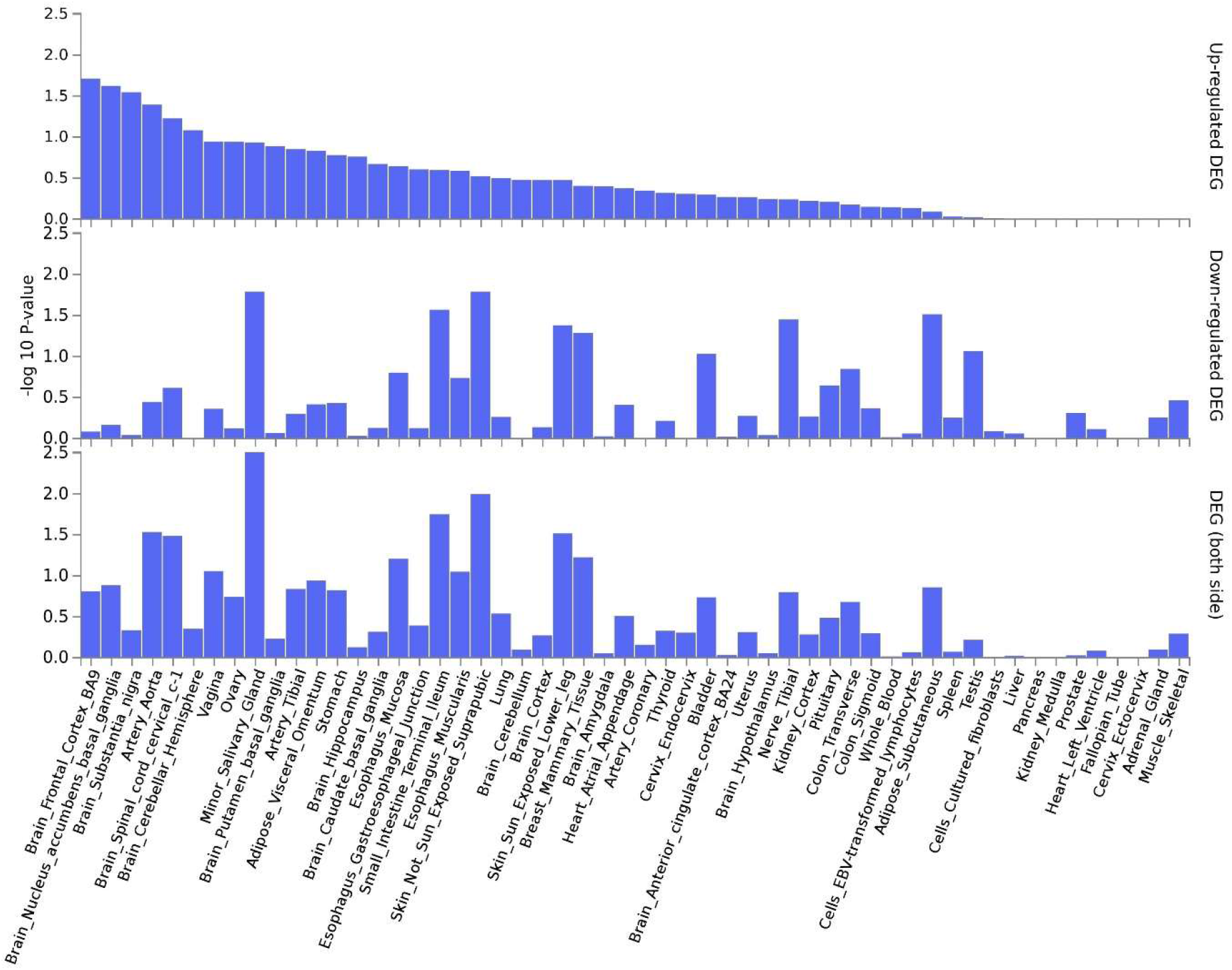
Up-regulated and down-regulated differentially expressed genes associated with having breakfast, generated by FUMA.

**Supplementary figure 10.**
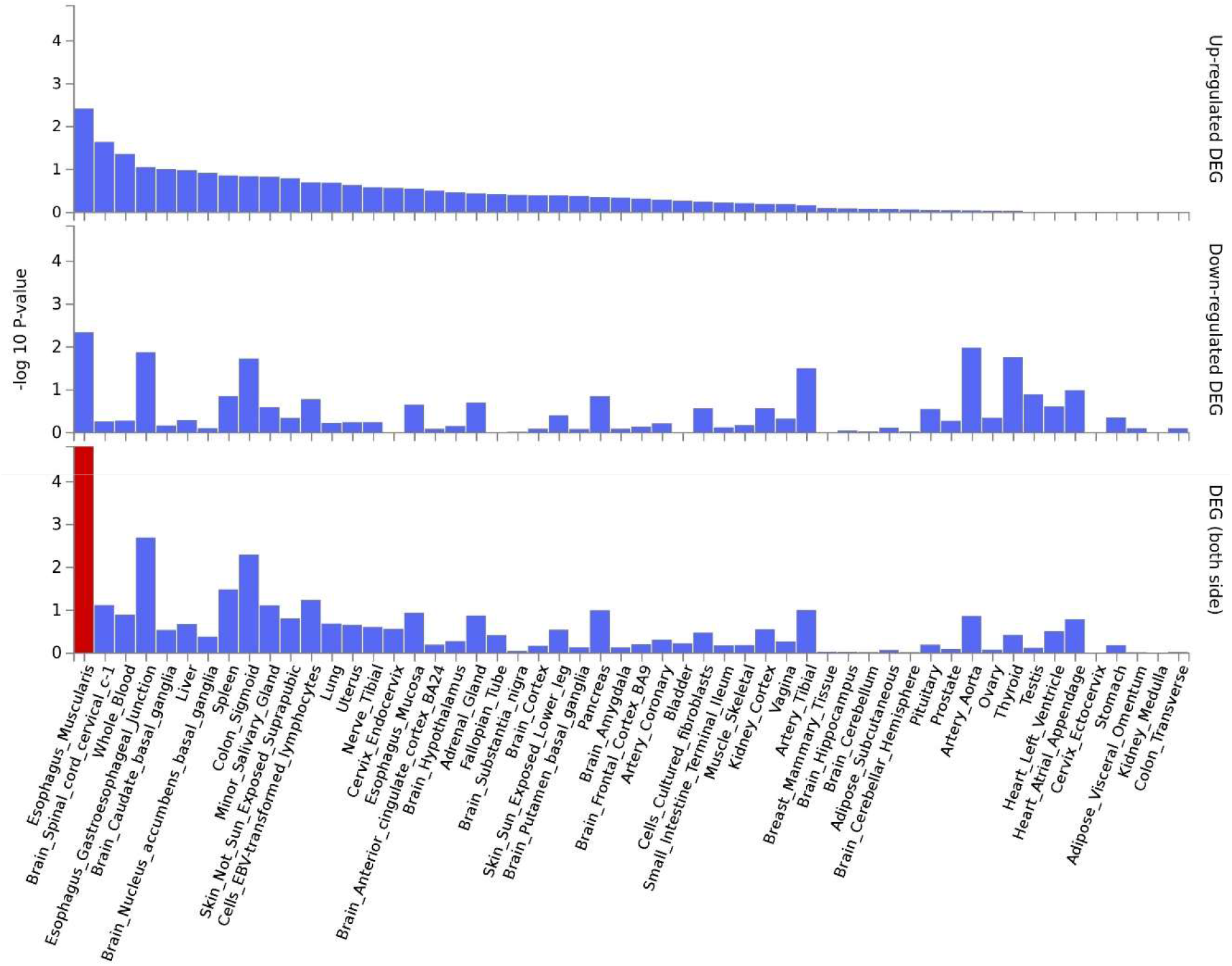
Up-regulated and down-regulated differentially expressed genes associated with having snacks, generated by FUMA.

**Supplementary table 5.** Genetic variants associated with measured confounders at a threshold of p=10^−4^ are excluded from the analysis.

| GRCh37/hg19 | rs number | Confounder | P-value |
| --- | --- | --- | --- |
| 16:2130686 | rs9936277 | household income | 1.48E-08 |
| 7:141886315 | rs141440087 | household income | 1.96E-07 |
| 14:95309911 | rs537485611 | household income | 6.75E-06 |
| 4:184499719 | rs74436000 | household income | 9.30E-05 |
| 2:215740962 | rs145316979 | father education | 3.59E-05 |

**Supplementary figure 11.**
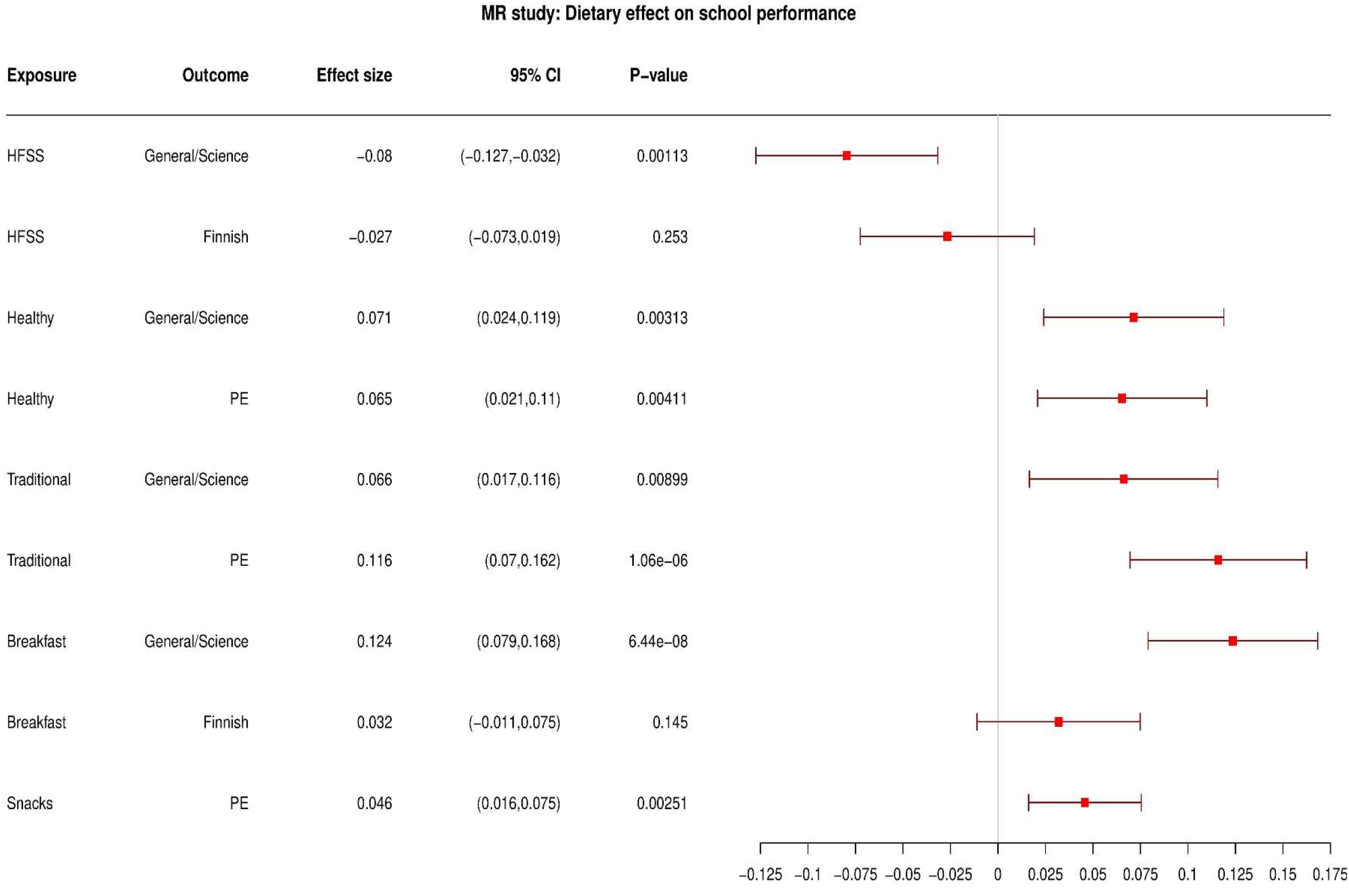
Results of one-sample mendelian randomisation: the effect of an additional weekly meal in dietary exposures on school performance outcomes. Genetic variants which are associated with measure confounders, below the threshold of P-value=10^−4^ are excluded from the analysis.

**Supplementary table 6.**
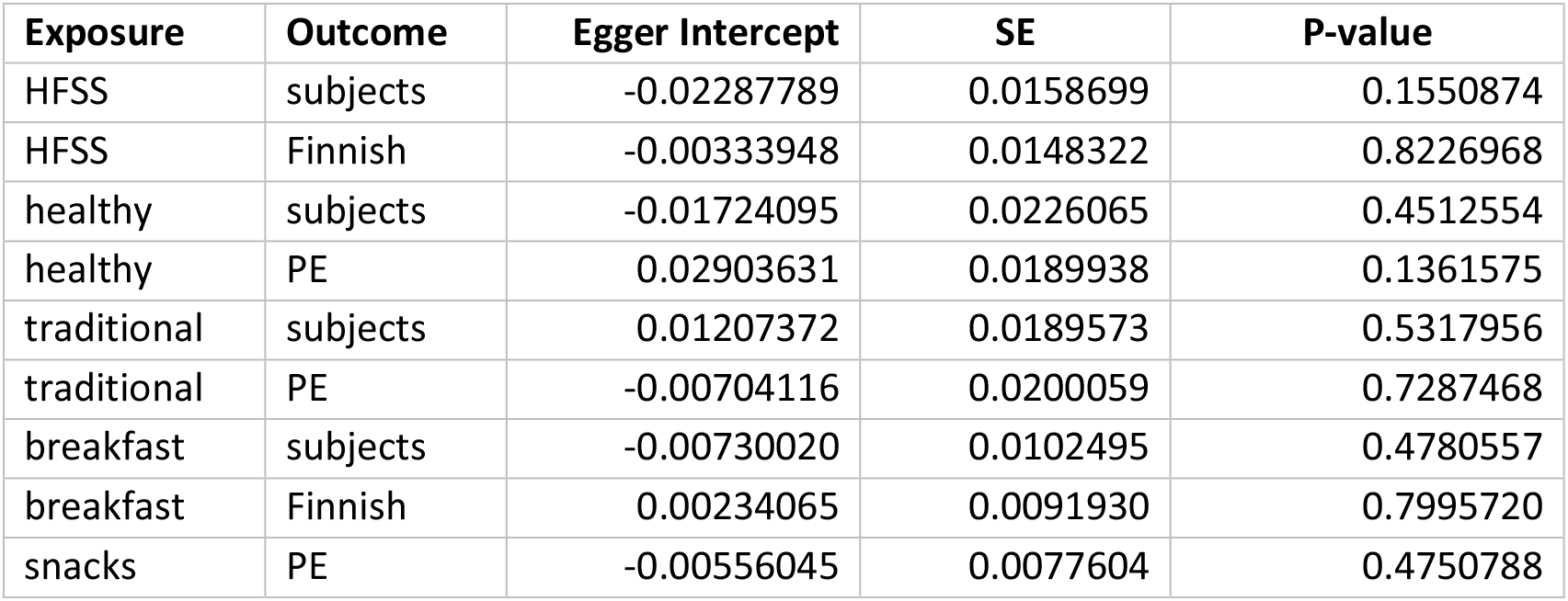
Mendelian randomisation – Egger intercept estimates.

**Supplementary table 7.**
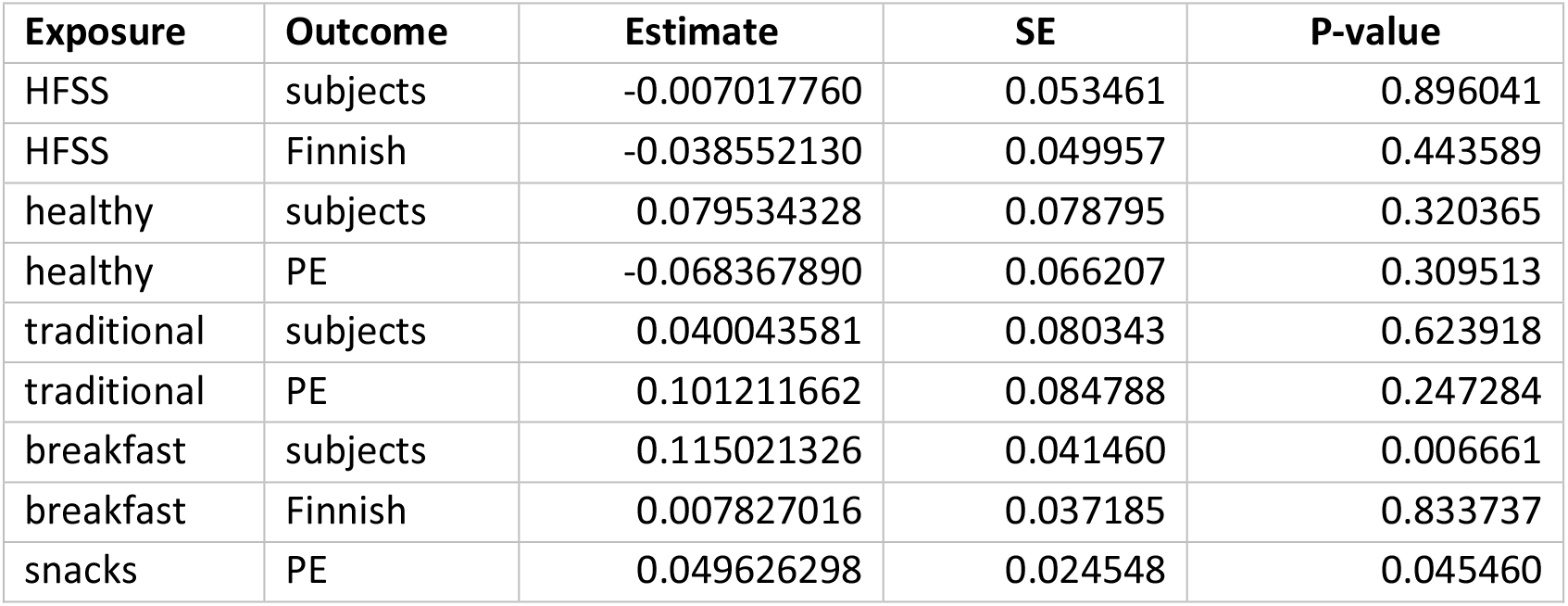
Mendelian randomisation – Egger effect size estimates.

| Exposure | Outcome | Estimate | SE | P-value |
| --- | --- | --- | --- | --- |
| HFSS | subjects | -0.007017760 | 0.053461 | 0.896041 |
| HFSS | Finnish | -0.038552130 | 0.049957 | 0.443589 |
| healthy | subjects | 0.079534328 | 0.078795 | 0.320365 |
| healthy | PE | -0.068367890 | 0.066207 | 0.309513 |
| traditional | subjects | 0.040043581 | 0.080343 | 0.623918 |
| traditional | PE | 0.101211662 | 0.084788 | 0.247284 |
| breakfast | subjects | 0.115021326 | 0.041460 | 0.006661 |
| breakfast | Finnish | 0.007827016 | 0.037185 | 0.833737 |
| snacks | PE | 0.049626298 | 0.024548 | 0.045460 |

**Supplementary table 8.** Weighted median mendelian randomisation results.

| Exposure | Outcome | Estimate | SE | P-value |
| --- | --- | --- | --- | --- |
| HFSS | subjects | -0.0484950 | 0.035168 | 0.167910 |
| HFSS | Finnish | -0.0243780 | 0.033617 | 0.468342 |
| healthy | subjects | 0.0277786 | 0.042531 | 0.513663 |
| healthy | PE | 0.0650143 | 0.039727 | 0.101729 |
| traditional | subjects | 0.1000436 | 0.054257 | 0.065201 |
| traditional | PE | 0.0712284 | 0.050550 | 0.158816 |
| breakfast | subjects | 0.0653719 | 0.023022 | 0.004518 |
| breakfast | Finnish | 0.0161014 | 0.014189 | 0.256478 |
| snacks | PE | 0.0154165 | 0.014458 | 0.286289 |

**Supplementary table 9.** Weighted mode mendelian randomisation results.

| **Exposure** | **Outcome** | **Estimate** | **SE** | **P-value** |
| --- | --- | --- | --- | --- |
| HFSS | subjects | 0.0432975 | 0.090051 | 0.632524 |
| HFSS | Finnish | 0.0346365 | 0.084102 | 0.682030 |
| healthy | subjects | 0.0485570 | 0.081229 | 0.554068 |
| healthy | PE | 0.0853619 | 0.078499 | 0.284725 |
| traditional | subjects | 0.1262579 | 0.105929 | 0.247248 |
| traditional | PE | 0.0826252 | 0.091188 | 0.375669 |
| breakfast | subjects | 0.0487474 | 0.057718 | 0.400449 |
| breakfast | Finnish | 0.0313670 | 0.039414 | 0.427701 |
| snacks | PE | -0.0078780 | 0.039112 | 0.840717 |

